# Kinobead Profiling Reveals Reprogramming of B-cell Receptor Signaling in Response to Therapy Within Primary Chronic Lymphocytic Leukemia Cells

**DOI:** 10.1101/841312

**Authors:** AJ Linley, LI Karydis, A Mondru, A D’Avola, S Cicconi, R Griffin, F Forconi, AR Pettitt, N Kalakonda, A Rawstron, P Hillmen, AJ Steele, DJ MacEwan, G Packham, IA Prior, JR Slupsky

## Abstract

Signaling via the B-cell receptor (BCR) is critical for driving CLL pathobiology, promoting both malignant cell survival and disease progression. However, understanding of this pathway is limited, particularly in relation to potential changes in response to therapy. Here, we describe a kinobead-based protocol, used in conjunction with mass-spectrometry to study surface-IgM signaling in primary CLL cells. We identified a ‘fingerprint’ of over 30 kinases which displayed unique, patientspecific response following sIgM stimulation. Matched analysis of CLL cells in samples taken from clinical trials showed that BCR-induced kinome responses altered between baseline and disease progression in patients who relapsed from chemoimmunotherapy. Moreover, adaptive changes to BCR signaling were also observed in CLL cells from clinical trial patients receiving ibrutinib; longitudinal profiling revealed increased signaling despite BTK inhibition. Collectively, these data comprise the first comprehensive investigation into BCR signaling response within CLL where kinobead profiling reveals unique evidence of adaptive reprogramming in response to therapy.

## INTRODUCTION

The accumulation and survival of chronic lymphocytic leukemia (CLL) cells is strikingly dependent on signaling from activated cell surface receptors.^1,2^ This applies particularly to the B-cell receptor (BCR) where antigen-dependent and –independent activity has been linked to disease development.^1,3-5^ For example, retained anti-IgM signaling capacity is associated with progressive disease,^6^ and kinase inhibitors (KI) targeted against BCR signaling pathways (especially the Bruton’s tyrosine kinase (BTK) inhibitor ibrutinib) can induce impressive clinical responses in patients with CLL and some subtypes of non-Hodgkin’s lymphoma.^7-9^ In addition to disease pathogenesis, altered signaling influences response to therapy. This is clearest in the context of acquired resistance to ibrutinib which is commonly associated with mutations of BTK itself or one of its downstream effectors, phospholipase Cg2 (PLCg2), where mutations within either result in production of proteins which retain activity but bind ibrutinib less avidly, or are hyper-responsive to upstream activating pathways, respectively.^10,11^

Kinase activation and inhibition in primary CLL cells has generally been analyzed by examining kinase-substrate phosphorylation using phospho-specific antibodies to identify modified amino-acids. While such an approach can provide important insight, interpretation of results can be challenging. For example, the activity of many kinases is influenced by multiple phosphorylation events which can be activating or inhibiting. Where phosphorylation at multiple sites on an individual kinase is detected, it can be difficult to determine whether these are on the same molecule, or whether subpopulations of kinases with distinct patterns of phosphorylation (and hence activity) co-exist. Furthermore, upstream kinases responsible for this phosphorylation can be inferred based on substrate specificity, but redundancy amongst kinases (where several kinases may be able to modify an individual phospho-acceptor site) limits the utility of this approach. Finally, changes in phosphorylation will reflect changes not only in kinase activity, but also their counteracting phosphatases.

Kinobeads provide a powerful tool to probe kinase function.^12-15^ In this approach cell lysates are incubated with beads coated with broad-specificity type 1 KI, allowing binding of a large proportion of the expressed kinome. Bound kinases can then be profiled using mass-spectrometry (MS). Kinobead-MS technology was initially deployed to determine the specificity profiles of KI since binding of KI to target kinases prevents their subsequent capture by the kinobeads.^16^ However, kinase capture can also be influenced by changes in the abundance of the kinases and, in particular, active site conformation. Many kinases exist in an auto-inhibited form and their activation requires conformational change. For example, BTK adopts multiple conformations allowing graded activation following cell stimulation,^17^ and ERK activation requires a MEK-dependent switch to an active conformation.^18^ Taking these considerations into account, kinobead technology has been used more recently to probe kinase activation in response to cell stimulation and, in particular, to reveal adaptive “rewiring” of kinase networks following exposure to KI.^12,19^

Here, we have developed the kinobead approach for analysis of malignant B-cells and used it for the first time to characterize signaling in primary CLL cells. We show kinobeads can be used to assess active site occupancy by kinase inhibitors, including in ibrutinib-treated patients.

## RESULTS

### Validation of kinobeads for the analysis of malignant B cells

We first examined the ability of kinobeads bearing different broad-specificity KI to capture kinases using the MEC-1 cell line. MEC-1 cells were derived from a CLL patient undergoing prolymphocytoid transformation^24^ and were selected because they have readily detectable levels of constitutively active kinases, including ERK1/2 and AKT (**Supplemental Figure 1A**). A schematic workflow for a typical experiment is shown in **Figure 1A**. Three separate experiments were performed.

**Figure 1:**
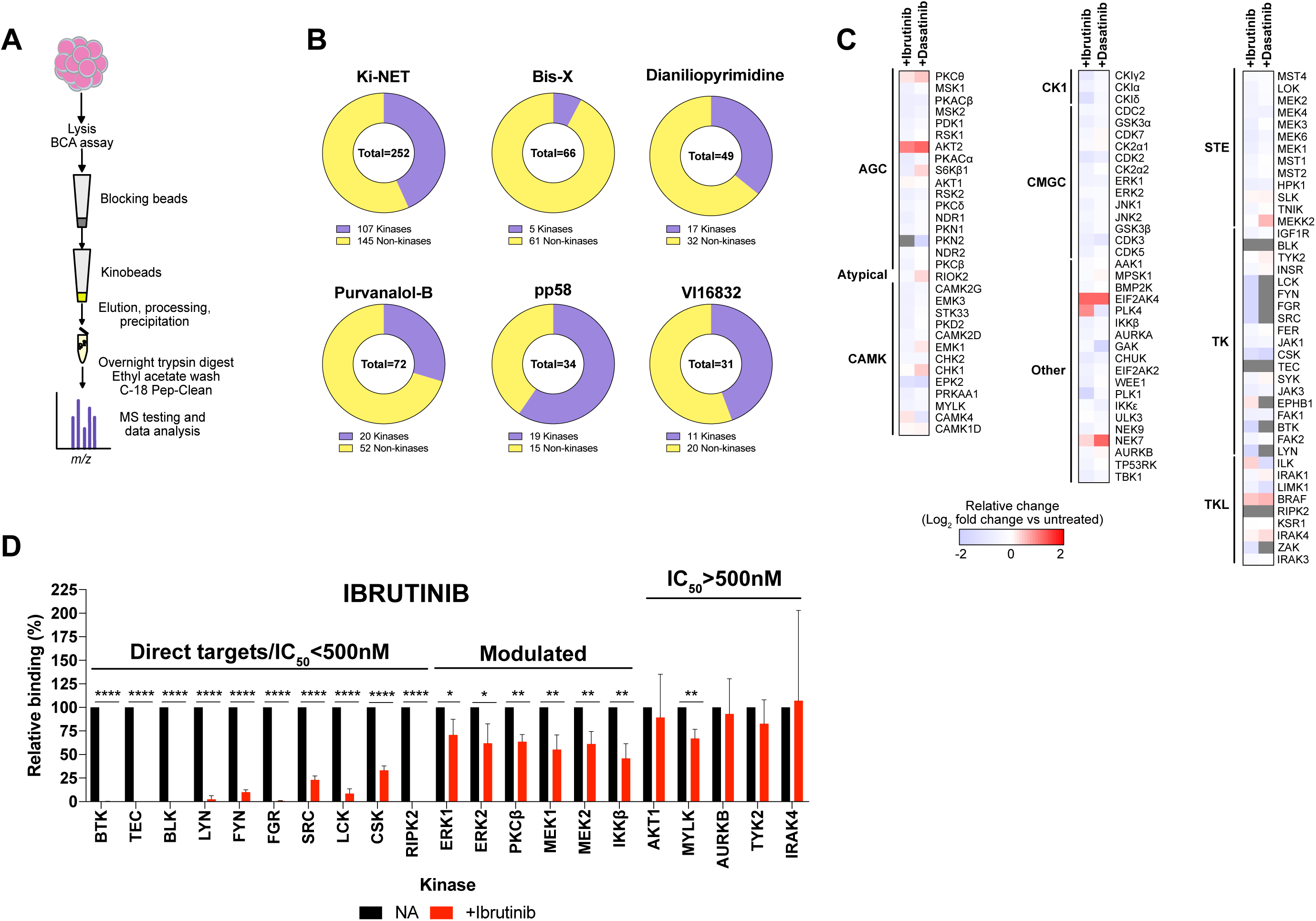
Kinobead analysis of MEC-1 cells. **A;** Schematic of the kinobead-MS used for this study. **B;** Pie graphs illustrating the numbers of kinase and non-kinase proteins isolated from MEC-1 cell lysates using 6 different kinobeads. **C;** MEC-1 cells were treated with ibrutinib or dasatinib (both at 500 nM) or left untreated as a control. After 60 minutes, cell lysates were prepared and kinase binding to Ki-NET beads was assessed using mass spectrometry. Kinases are grouped by subfamilies. Heat map shows mean change (log_2_) in kinase binding for ibrutinib or dasatinib-treated cells relative to control cells for the core 107 kinases derived from three independent experiments. **D;** Graph showing results for ibrutinib for selected kinases, including direct targets (i.e., inhibited by ibrutinib in *in vitro* assays with IC_50_ <500 nM), kinases which are modulated by ibrutinib but are not direct targets, and kinases which are unaffected by ibrutinib. Graph shows mean (±error) binding normalized to control cells (set to 100%) with statistical significance of differences indicated (Student’s ttest; *=P<0.05; **=P<0.01; ****=P<0.0001).

We found that the most effective kinobeads for kinase capture were those bearing KiNET (CTx-0294885). These isolated 132 kinases in total (i.e. identified in at least one of the experiments performed), with a core set of 107 kinases being detected in all three experiments (**Figure 1B, C; Supplementary List**). This core set of captured kinases included representatives from all kinase sub-families and well-characterized BCR signalosome components, including LYN, SYK and BTK (**Figure 1C**). By contrast, kinobeads bearing bisindolylmaleimide-X (Bis-X), dianillopyrimidine, purvalanol-B, pp58 or VI16832 captured substantially fewer kinases (**Figure 1B**). Each kinobead also captured various non-kinases and, in some instances, the number of non-kinase targets captured substantially outweighed the number of kinases (e.g. purvalanol-B). Identified non-kinases may reflect direct capture of non-kinase targets or indirect capture of kinase-associated proteins. Importantly, ∼70% of the unique kinases identified using any of the 6 kinobeads were captured using Ki-NET only (**Supplementary Figure 1B**) and we therefore focused on Ki-NET kinobeads for further studies. Comparison of intensities determined by MS analysis for the core 107 kinases detected using Ki-NET found a highly significant correlation between the biological repeats for control and inhibitor-treated cells, thereby confirming the reproducibility of the technique (**Supplementary Figure 1C**).

We used Ki-NET kinobeads to investigate the effects of two tyrosine KI on the core set of 107 kinases captured from MEC-1 cells (**Figure 1C**). Ibrutinib and dasatinib are considered primarily as inhibitors of BTK and ABL, respectively, although both drugs have numerous off target effects, some of which are shared.^25^ Analysis was performed after 60 minutes of drug exposure; a time point at which we would not expect to observe substantial changes in the steady state expression of kinases. Thus, changes in kinase capture are likely to reflect both direct target engagement (thereby blocking kinobead binding) and secondary effects on downstream kinases.

The most striking effects of ibrutinib and dasatinib on kinase capture were observed within the TK and TK-like (TKL) families and appeared to be due to occupancy of direct targets (**Figure 1C**). Thus, BTK capture was reduced by >99% in cells treated with ibrutinib (**Figure 1C, D)**. BTK capture was also essentially ablated in cells exposed to dasatinib **(Figure 1C)**, which is also known to inhibit BTK.^26^ Other kinases with substantially reduced capture in drug treated cells included LCK, FYN, LYN, FGR, SRC, CSK, TEC, BLK and RIPK2, which are all known to be directly inhibited by both ibrutinib and dasatinib.^25-28^

We also observed more modest reductions in capture of other kinases including ERK1 and 2, PKCb and IKKb **(Figure 1D)** which are not known as direct targets of ibrutinib/dasatinib but are likely to be modulated as a response to upstream effects. Reduced capture of MEK1 and MEK2 was consistent with the observed reduction of phosphorylation of the MEK1/2 substrates ERK1/2 detected by immunoblotting (**Supplementary Figure 1**). Thus, kinobeads reveal both the direct occupancy of kinase active site by drug, as well as indirect modulation of active site availability of non-target kinases.

### Kinobead analysis of primary CLL cells

#### Response of CLL kinome to sIgM stimulation

We next performed Ki-NET kinobead analysis of primary samples from a cohort of 40 CLL patients. Profiling was performed using cells treated either with control antibody or with anti-IgM to allow us comparison between the “basal” signature in the absence of exogenous stimulation and that in response to sIgM stimulation, respectively. Analysis was performed at 5 minutes following addition of anti-IgM (or control antibody) since previous analyses showed that this was suitable time point for analysis of both proximal (e.g. LYN, SYK) and distal (e.g. ERK1/2) signaling responses in CLL cells. To avoid artefacts associated with extended cell manipulations we did not purify malignant cells before stimulation/analysis. The average proportion of malignant cells in the samples used was 86%, no T-cell-specific kinases, such as ITK, were detected at any time and during our analyses, suggesting that T-cell contamination minimally affected this assay. In all cases, cell viability was ^3^ 90%. Initial experiments identified 104 kinases from lysates of both control and IgM-stimulated CLL cells. Comparison of this signature with that isolated from MEC-1 cells showed, as expected, considerable overlap where 74% of these 104 kinases were common, demonstrating that the kinome fingerprint discovered with MEC-1 can largely be replicated using primary CLL B-cells. Furthermore, the ability of kinobeads to capture kinases appeared independent of their expression level; Nanostring analysis of kinase and pseudokinase mRNA levels in CLL cells (**Supplementary Figure 2A; Supplementary List**) showed that the major kinase subfamilies identified by both techniques had proportionately similar representation and that individual kinases could be identified by the kinobeads across a wide range of expression.

Analysis of the entire cohort showed that capture of some kinases was relatively consistent across all CLL samples, whilst others were more variably detected. As such, all 104 kinases described were detected in only half of the cohort of samples we analyzed. Therefore, we determined a more restricted, core fingerprint of 32 kinases that were identified in lysates from at least 75% of patient samples (**Supplementary List**) Importantly, this core fingerprint included key kinases involved in BCR signaling, indicating that kinome profiling of CLL cells is able to capture and detect change to the central elements of this pathway.

To probe the reproducibility of our approach using primary cells, biological repeats were carried out. Comparison of MS intensities for control and anti-IgM-treated cells revealed significant correlation across biological repeats (**Supplementary Figure 2B**), further confirming the robust nature of the kinobead technique. To the best of our knowledge, this represents the most comprehensive signaling profile of primary CLL cells. Thus, we found, in agreement with the known heterogeneity of sIgM signaling capacity in CLL cells,^3,29^ that there was substantial variation in the response to anti-IgM between samples analyzed using kinobeads (**Figure 2; Supplementary Figure 2**). Some samples showed relatively few changes in our identified core fingerprint of kinases following anti-IgM-stimulation, whereas others showed much more change. Assembly of kinome response into heatmaps revealed how each patient demonstrated a unique profile of response. These responses were not clearly related to prognostic indicators such as *IGHV* mutation status, Binet staging or karyotype (**Supplementary Figure 2C**). We therefore focused on analyzing the kinome fingerprint in terms of signaling indicators, specifically capacity to induce intracellular calcium flux (iCa^2+^) and sIgM expression (**Figure 2A**). For the former, we compared patients based on their *in vitro* signaling capacity (Signaler vs Non-Signaler) as previously described.^21^ Importantly, we identified increased capture of SYK and BTK in Signaler compared to Non-Signaler patients (**Supplementary Figure 2D**). These kinases are key kinases in the BCR signaling pathway and both are more highly represented in samples which induce iCa^2+^ flux in response to sIgM crosslinking than in samples which do not. These data suggest that kinobeads measure canonical BCR signaling in CLL cells that is in line with traditional methods, but also provides information on additional pathways engaged in response to activation of sIgM.

**Figure 2:**
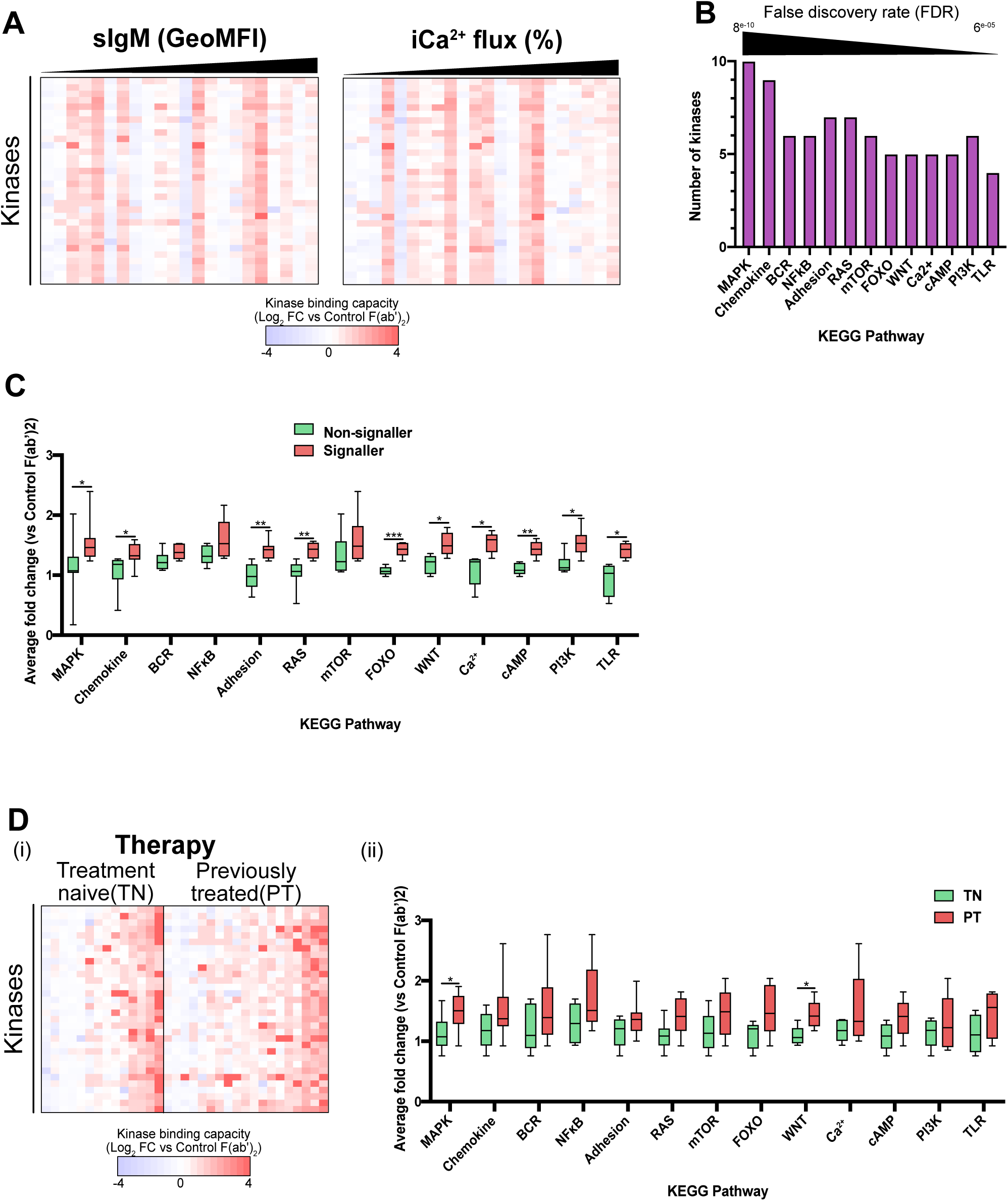
Kinobead profiling of primary CLL cells. Cell lysates were prepared from CLL samples (n=40) incubated with control F(ab’)2 antibody (no stimulation) or anti-IgM. Kinase binding was assessed using Ki-NET beads and mass spectrometry. **A;** Heatmaps showing signature of 32 kinases, illustrating relative changes in kinase binding (log2 fold change vs control cells) in response to sIgM stimulation for each patient. Each patient (columns) was stratified based on either sIgM expression or the proportion of malignant cells capable of intracellular Ca^2+^ flux (iCa^2+^ flux) **B;** Graph derived from STRING analysis of the 32 kinase fingerprint, illustrating the KEGG pathways identified by STRING analysis and the number of kinases from the fingerprint that are involved in each pathway. Pathways are ordered based on false discovery rate (FDR). **C;** Graph to show changes in identified KEGG pathways in Signaler patients compared Non-Signaler patients in response to anti-IgM stimulation. Y-axis shows on average change in abundance for detected kinases within each KEGG pathway (Student’s t-test; *=P<0.05; **=P<0.01; ***=P<0.001). **D**; Heatmap showing relative changes to kinome capture between Signaler patients stratified according to receipt of chemoimmunotherapy. Patients were classed as being treatment naïve (TN) or previously-treated (PT) patients. **E**; Comparison of changes to KEGG pathways associated with our kinome signature between TN and PT patients.

The individual kinome fingerprints associated with CLL cells from each patient contained a diverse range and number of kinases. This implied the potential for a wide range of signaling networks which could be affected following stimulation of sIgM. To explore this, we next probed the nature of these additional pathways further by applying STRING analysis to the core fingerprint of 32 kinases that were captured by kinobeads. Using a strict false discovery rate (FDR) (<0.0001) we identified 13 KEGG pathways associated with this kinome fingerprint (**Figure 2B**), highlighting the shared roles/function of the individual kinases. Reassuringly, we not only identified the BCR pathway as a key highlight, but also others known to be associated with CLL cell signaling, the MAPK, mTOR, TLR and NFkB pathways. Interestingly, other pathways such as those associated with FoxO, RAS, cAMP and WNT, that have been less studied in relation to CLL cell biology were also identified. Using this approach, we found that cells from Signaler patients demonstrated significant increases in 10/13 (77%) of these KEGG pathways compared to Non-Signaler patients (**Figure 2C**). These findings, taken together with our observations of signaling variability in CLL cells from different patients, suggest the potential of a significantly wider signaling network associated with sIgM activation that involves novel influences that may be individual to the environment or dominant clone of CLL cells within each patient.

#### Therapy influences sIgM response in primary CLL

The heterogeneity recognized in CLL patients applies not only to their biology and clinical presentation, but also to receipt of therapy. Studies in other tumor types have described treatment-related alterations in signaling. To explore this possibility in CLL, we stratified Signaler patients within our cohort based upon receipt of chemoimmunotherapy (CIT). Intriguingly, when responses were stratified in this manner, the scale of responses shown by treatment naïve (TN) patients was remarkably similar to those who had previously received treatment (PT) (**Figure 2Di**). Comparison of TN and PT patients found the latter group displayed increased changes in all 13 KEGG pathways, which in the case of MAPK and WNT were significantly different (p=<0.05; Student’s T test) (**Figure 2Dii**). Importantly, these alterations of the kinome were not due to differential expression because comparison of kinome mRNA levels found a highly significant level of correlation (P=<0.0001; **Supplementary Figure 2E**), nor were they due to changes in phosphatase activity (**Supplementary Figure 3**) because we were unable to observe any differences in the expression of key B-cell phosphatases ^30-34^. Taken together, these data suggested the presence of signaling patterns resulting from changes to kinase activation based upon receipt of therapy.

#### Kinobeads can be employed to recognize reprogramming of BCR signaling

Our data indicating therapy-induced alterations was reminiscent of modulation of the BCR signaling network induced through the addition of various factors. A key example of this can be seen through the modulation of BCR signaling by factors present in the CLL microenvironment. A strong candidate is interleukin 4 (IL4), which, when used to pre-treat primary CLL cells in vitro, potentiates sIgM signaling.^22^ We therefore sought to examine if we could use kinobead-MS to determine changes brought about through IL4 in a group of 5 Signaler patients. Cells were pre-treated with IL4 as described previously^22^ or left untreated prior to anti-IgM stimulation and lysate collection (**FIGURE 3A**). Kinobead profiling revealed that conditioning of CLL cells by IL4 resulted in alterations to the kinome fingerprint for each patient (**FIGURE 3B**), therefore suggesting adaptation of the signaling response. Moreover, analysis of the KEGG pathways outlined above found that 8/13 (62%) of these were significantly up regulated when cells were treated with both IL4 and anti-IgM (**FIGURE 3C**). These data demonstrate that kinome profiling can detect reprogramming of BCR signaling by IL4. Taken together with our data for the main patient cohort, this suggested that not only does the BCR response involve a wider signaling network than previously appreciated, but that this response can be reprogrammed by treating CLL cells with agents such as IL4.

**Figure 3:**
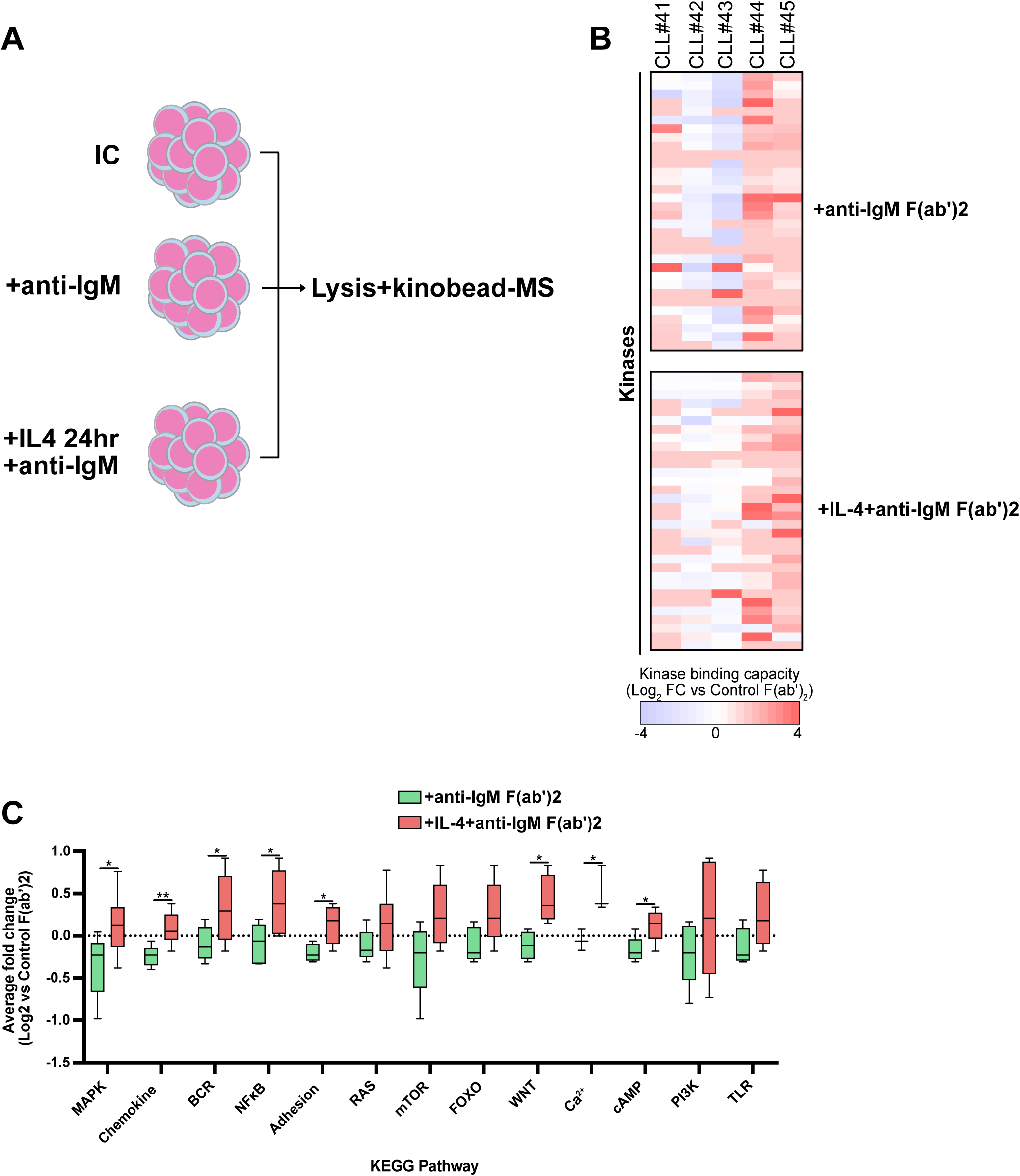
Kinobead-MS can detect reprogramming of sIgM signaling induced through exposure to IL4. **A**; Schematic cartoon illustrating cell conditions/treatments prior to protein lysis and incorporation into the kinobead assay. **B**; Heatmaps showing relative changes to kinome signature in a group of 5 CLL patients stimulated with anti-IgM (±overnight pre-treatment with IL4). **C**; Comparison of changes to KEGG pathways (log2 scale) within these 5 patients in response to anti-IgM alone or a combination of IL4 and anti-IgM.

### Detection of intra-patient signal reprogramming influenced by therapy

#### Chemoimmunotherapy

To further investigate the role of possible influence of therapy on sIgM-induced signaling, we next performed matched sample analysis. In the first instance, this involved samples from patients (n=10) who had been treated with FCR (fludarabine, cyclophosphamide, Rituximab) within the ARCTIC/ADMIRE clinical trials. Kinobead-MS was carried out on the baseline (BL) samples associated with these patients prior to treatment and then again on the matched sample taken at disease progression (DP). The unique, patient-specific nature of kinome responses upon antiIgM stimulation was once again apparent in the BL samples. Remarkably, this kinome response completely changed in the matched patient sample taken at DP (**Figure 4A**), suggesting adaptation of BCR signaling response related to time and therapy. Comparison of relative changes to the 13 KEGG pathways we previously identified between timepoints found a number of pathways which were significantly upregulated at progression compared to baseline (**Figure 4B**). Interestingly, while the BCR pathway did not appear to alter between these timepoints, the MAPK pathway, along with RAS, mTOR, Chemokine, Adhesion, FOXO and WNT were all significantly increased. A number of kinases (e.g. PKCb, ERK2, JNK2, RSK1/2) showed significantly greater representation within kinome profiles of DP samples than in BL samples (**Figure 4C**). These kinases are common to the up-regulated signaling pathways and have demonstrated relation to CLL pathogenesis,^1,3^ suggesting that they are potential drivers of pathway change. Other kinases also exhibited significant fold increase in change (NDR1, TBK1, MST1) within the kinome profile of DP patients (**Figure 4C**). The role of these kinases is not described either within the BCR signaling pathway, nor in CLL pathogenesis, indicating novel involvement of other potential drivers of BCR signaling pathway change. Collectively, these data suggest a novel finding that BCR signaling rewiring within the adaptive response of malignant cells from CLL patients when they relapse from chemoimmunotherapy.

**Figure 4:**
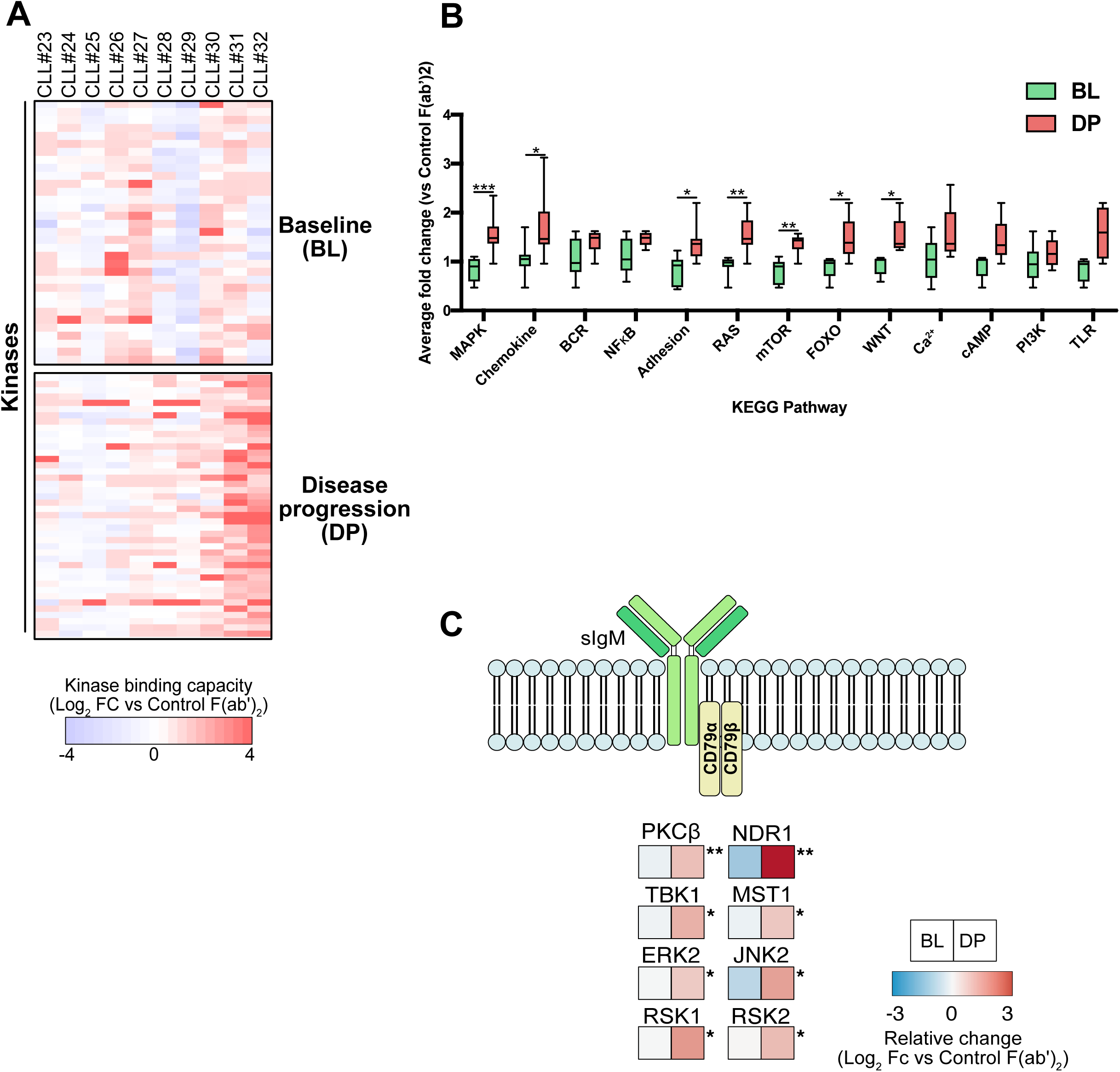
Matched kinobead profiling of sIgM response in ARCTIC/ADMIRE patients. **A;** Heatmap comparing sIgM activation kinome fingerprints for ARCTIC/ADMIRE trial patients (n=10) using their baseline (pretherapy) sample and their disease progression (DP) sample. **B;** Graph comparing changes in KEGG pathways between BL and DP patients. **C;** Signaling atlas for kinase nodes found to be most significantly changed within the studies KEGG pathways (log2 scale) upon activation of sIgM.

#### Ibrutinib

Small molecule agents have become increasingly prevalent as therapy options for CLL, the most evident example of this being ibrutinib which is becoming a standard of care in the treatment of this disease. By targeting BTK, ibrutinib blocks the function of a key kinase within the BCR pathway to limit cell proliferation. However, about a third of CLL patients relapse on ibrutinib therapy, the basis of which not necessarily related to resistance mutations in either BTK or PLCg2. Considering our findings using patient samples from the ARCTIC/ADMIRE trials, we next sought to investigate whether ibrutinib treatment caused similar reprogramming of sIgM signaling using samples derived from patients recruited to the IcICLLe clinical trial.

We performed longitudinal profiling on 4 patient samples to compare baseline (preibrutinib) with those taken 1 month following initiation of ibrutinib (**Figure 5A**). In this instance, certain kinases from the core fingerprint could not be detected in both samples, meaning we identified a signature of 26 kinases which was common between both time points (**Supplementary List**). Interestingly, this signature altered dramatically, displaying a marked increase in intensity in samples from patients receiving ibrutinib (**Figure 5A**). This increase in intensity was not related sample by sample to lymphocyte count but could have relation to sIgM expression on the malignant cells in a way that is in line with previous work^35^ showing that circulating CLL cells from patients undergoing ibrutinib therapy demonstrate increased sIgM levels. (**Supplementary Figure 4A**). Importantly, in the ibrutinib-treated samples there was a loss of BTK and BLK from the fingerprint. This loss of BTK binding to KiNET was confirmed using immunoblotting of lysates derived from cells from patients undergoing ibrutinib therapy within our clinics (**Supplementary Figure 4B**), illustrating the *in vivo* action of ibrutinib to block access of BTK to the Ki-NET beads.

**Figure 5:**
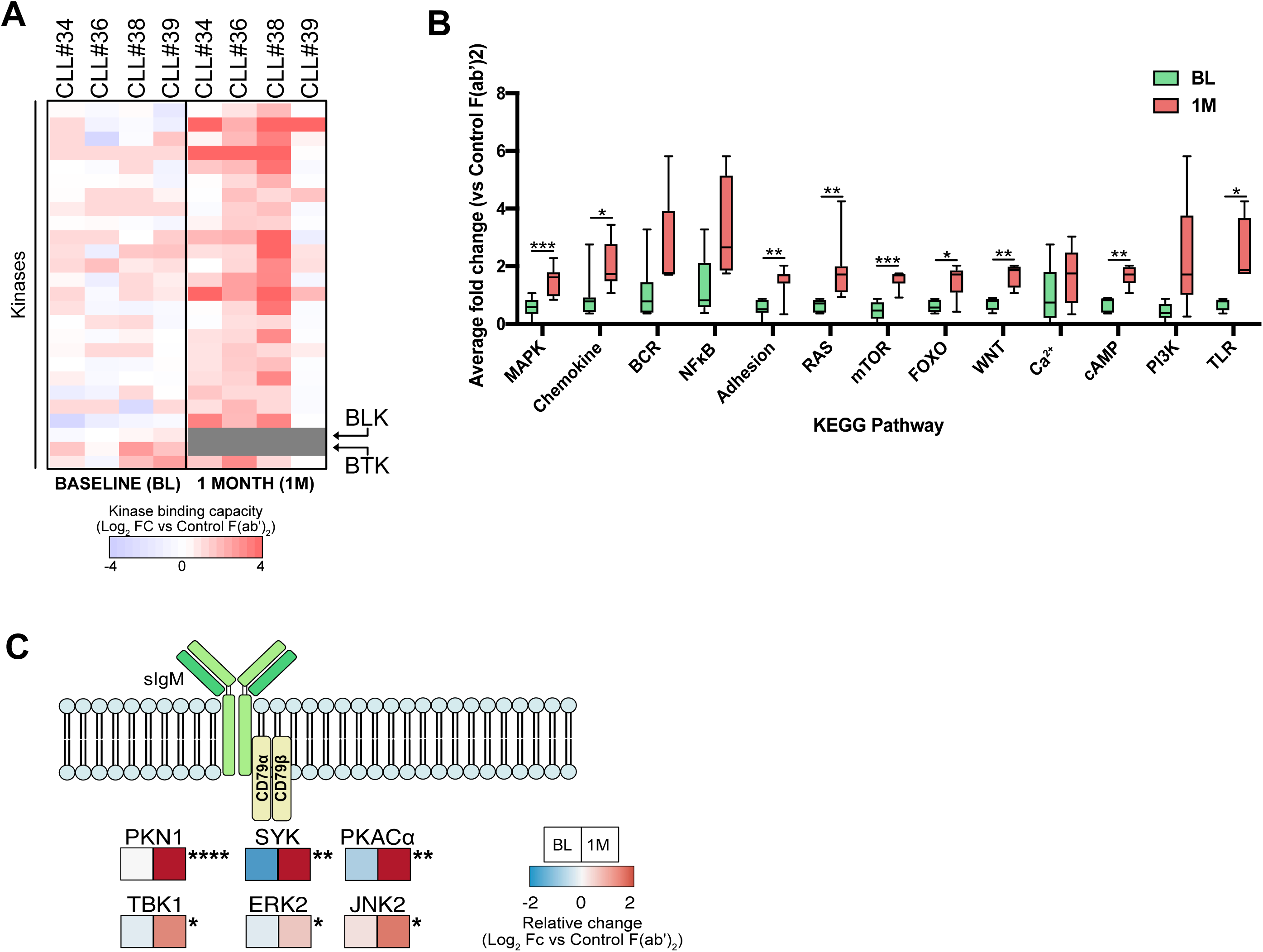
Longitudinal kinobead profiling of sIgM response in IciCLLe patients. **A;** Heatmap comparing sIgM activation kinome fingerprints for IcICLLe trial patients (n=4) using their baseline (pre-therapy) sample and samples taken 1 month following commencement of ibrutinib therapy. Grey boxes denote loss of kinase binding. **B;** Graph comparing changes in KEGG pathways between BL and 1month (1M) samples. **C;** Signaling atlas for most significantly altered kinases illustrating the relative change (log2 scale) upon activation of sIgM.

This is likely to also apply to BLK, as previous work has shown ibrutinib can target both of these kinases.^27^

Further analysis of the kinome signature intensity in ibrutinib-treated patient samples showed it was accompanied by increases in KEGG pathways in a similar manner as observed in the FCR-treated patient samples (**Figure 5B**). Overall, the KEGG pathways that were significantly altered in response to ibrutinib were the same as what was observed in response to FCR, but with the addition of cAMP and TLR pathways. Interestingly and although not significant, there was a marked increase in BCR pathway signaling despite the loss of BTK due to ibrutinib. Such increase in signaling is likely due to the significant increased representation of SYK, ERK2 and JNK2 within the kinome profiles isolated from the ibrutinib-treated samples (**Figure 5C**). Alternatively, other potential drivers of signaling are observed, notably PKN1, TBK1 and PKACα (**Figure 5C**). These data suggest a compensatory mechanism of signal pathway reprogramming in CLL cells taken from patients exposed to ibrutinib therapy that is activated within a month of receiving treatment.

## DISCUSSION

In the current study we employed a kinobead-based approach to examine BCR signaling within primary CLL cells. The results we generate are, to the best of our knowledge, the most comprehensive profiling of signaling mediators within the BCR signaling pathway in these cells. An important and novel finding we make is adaptation in sIgM signaling brought about through receipt of therapy. Thus, we show that CLL cells from patients who had received either traditional CIT or ibrutinib change their response to BCR engagement. Our analyses further suggest new drivers of signaling within the BCR pathway become involved and are specific to the therapy received. Our study therefore creates new understanding of the role BCR signaling plays within the natural history of CLL.

The most common method for investigating signaling in CLL involves the use of antibodies to screen restricted numbers of kinases and phosphoprotein targets using immunoblotting or flow cytometry. While informative, these methods lack the resolution required to observe widespread changes to signaling mediators within cells in response to various conditions. Alternatively, mass-spectrometry (MS) is an emerging technology in terms of analyzing signaling in CLL cells, used so far only to provide a global view of protein expression in relation to investigate malignant cell biology and its relation to disease severity.^36-38^ In our approach we used a kinobeadbased protocol to isolate signaling proteins that could then be identified by MS to ultimately examine changes to sIgM signaling in CLL cells. This method reproducibly isolated a profile of over 100 kinases from MEC-1 and primary CLL cells representing good coverage of all major kinase subfamilies expressed in these cells, because of the high correlation between kinases identified by mRNA expression and by kinobead profiling. Thus, kinobeads were able to detect a substantial proportion of the kinome in the malignant B-cells from patients with CLL, a finding that is consistent with kinome profiling studies on other cell types.^12-15^

Our analysis of response to sIgM engagement in primary CLL cells showed that the fingerprint of activated kinases from individual samples varied widely. Such signaling heterogeneity will be related to the acknowledged variability of CLL cell response to BCR engagement ^3, 21^ and our data provides new insight into this phenomenon through identification of 13 KEGG pathways that show change in the kinome signatures between Signaler and Non-Signaler cells. While some of these pathways are readily recognized as being associated with the pathobiology of CLL, some are less well known and reveal potential new drivers of signaling within the BCR pathway. Our data show that these drivers are particularly important for CLL cell adaptation of BCR signaling in response to therapy. We applied kinome profiling to matched patient samples taken at baseline and then again following disease progression from the ARCTIC and ADMIRE clinical trials testing the efficacy of fludarabine, cyclophosphamide, Rituximab (FCR) treatment. We found that 6 of the 13 identified KEGG pathways showed significant change between baseline and disease progression and implicated the kinases listed in Figure 4C as drivers of this change. In a similar way, analysis of matched samples from the IcICLLe clinical trial testing the efficacy of ibrutinib revealed that 9 of the 13 identified KEGG pathways had significant change between samples taken at baseline and after 1 month of therapy. Importantly, the drivers responsible for this change in response to ibrutinib therapy, listed in Figure 5C, are different from those responsible for the changes observed in samples taken from FCRtreated patients. Together, these data suggest reprogramming of the BCR signaling pathway can take place in CLL cells, a notion that is strongly supported by the reprogramming effect IL4 has on BCR signaling in CLL cells demonstrated within a study of BCR-induced Ca^2+^-flux^22^ and from our kinome profiling experiments shown in Figure 3. Such reprogramming might be an expected response of CLL cells in patients on ibrutinib therapy, similar reprogramming was observed in (AZD6244)treated triple negative breast cancer cells where kinobead analysis has demonstrated adaptive activation of receptor tyrosine kinases in response to MEK inhibition,^12^ and in other malignant cell systems and small molecule agents.^39,40^ However, this is the first time that signal pathway reprogramming is described as a potential resistance mechanism for cancer cells and cytotoxic therapies, and is particularly exciting for our understanding of CLL and the role BCR plays in this disease.

An intriguing aspect of our analysis is the identification of kinase nodes that could act to drive the adaptive potential of sIgM signaling in response to therapy. PKCb, ERK2, JNK2, RSK1/2 and TBK1 are well described mediators/effectors of CLL signaling involving the MAPK and NFkB networks that are central for malignant cell proliferation and survival in this disease.^1-3^ Roles for NDR1 (STK38) and MST1 (STK4) have yet to be described in CLL, but these key regulators of the Hippo pathway are implicated in BCR signaling where they can affect activation of BTK.^41^ In a similar way the functions of HPK1 (MAP4K1) and PKN1 in CLL cells are also unknown but is implied in studies showing that the former is involved in BCR signal rewiring within germinal centers,^42^ and that the latter is involved in lymphocyte migration.^43^ One possible explanation for our observations is reduced phosphatase activity, which could influence a large number of kinases. However, we were unable to identify a clear relationship between expression of a panel of phosphatases/inhibitory receptors and treatment status to support this concept.

A unique feature of kinome profiling is the ability to analyze drug occupancy in primary CLL cells from patients receiving ibrutinib therapy. Ibrutinib covalently modifies BTK at Cys^481^ and is therefore irreversibly bound to this protein where it blocks access to ATP and compounds that mimic ATP. We show that the ability of the inhibitor beads to isolate BTK and other tyrosine kinase where the Cys residue is in a relevant position (i.e. BLK) to be modified by ibrutinib, is eliminated in samples taken from patients on this type of therapy. This provides a novel molecular tool to investigate drug dosing in patients whereby the minimal dose of ibrutinib, or similar drug, can be determined that still affects BTK function. Such knowledge is useful in the quest to reduce/remove unwanted side effects.

Treatment with KI is becoming ever more prevalent to target cancer. This is especially evident with ibrutinib, which has proven to be a highly effective agent for the treatment of B-cell tumors including CLL. Although resistance to this agent has emerged, its overwhelming beneficial impact in terms of survival when compared to CIT^44^ has led ibrutinib to become FDA-approved as front-line therapy for treating patients in association with Rituximab. Our data highlight the ability of patients receiving CIT to experience changes to the kinome response, which has implications for their disease course and influence of second line treatments. Future work would involve examining the potential for CIT-induced signaling changes to possibly influence efficacy of ibrutinib. Our study shows that rewiring of signaling needs now to be recognized as a potential major issue to the treatment of CLL patients. We propose these findings act as basis to drive for personalized medicine or at very least a signal-omics approach to investigating effects of therapy in CLL.

## MATERIALS AND METHODS

### Cells

Primary cells from patients diagnosed with CLL according to iwCLL-2008 criteria were obtained from patients attending clinics at Royal Liverpool and Broadgreen Hospitals at the University of Southampton Hospital trust in the Mature lymphoid malignancies Observational Study (NIHR/UKCRN Portfolio ID: 31076). All patients provided informed consent. Samples were provided as heparinized whole blood from which peripheral blood mononuclear cells (PBMC) were isolated using Lymphoprep (Axis Shield, Oslo, Norway) and cryopreserved as previously described.^20^ Alternatively, primary cells from patients recruited to the ARCTIC/ADMIRE (ISRCTN16544962) and IcICLLe (ISRCTN12695354) clinical trials were obtained from the UK CLL Clinical Trials BioBank (Liverpool, UK). In all cases, cells were recovered as outlined before,^20^ counted and diluted to 1×10^7^/ml using pre-warmed complete medium (RPMI1640+Glutamax (Gibco) supplemented with 10% (v/v) fetal calf serum (FCS) and 1% (v/v) Pen/Strep). The CLL-derived cell line MEC-1 was cultured in complete medium. Cell line identity was confirmed using STR analysis (Powerplex 16 System, Promega) and the absence of mycoplasma was confirmed using Mycoplasma PCR detection kit (Applied Biological Materials).

### Characterization of primary CLL cells

Cells from non-trial patients underwent cell surface phenotyping using anti-CD19, anti-CD5 (both Biolegend, Cambridge, UK), anti-IgM (Dako, Ely, UK) and appropriate isotype control antibodies, and using an Attune NXT flow cytometer (Thermo Fisher Scientific, Loughborough, UK) FACScan or a FACS Canto (both BD Biosciences, Oxford, UK). Signaling capacity was determined by measuring the proportion of malignant cells that were able to flux intracellular calcium (iCa^2+^) following treatment with F(ab’)_2_ anti-IgM, as described previously^21^ using a FACScan or FACS Calibur instrument (BD Biosciences). All flow cytometry data were analyzed using FlowJo (TreeStar, Oregon, USA).

### Treatments

Primary CLL samples were treated with 20 µg/ml anti-IgM F(ab’)_2_ or control F(ab’)_2_ (both Southern Biotech, Cambridge Biosciences, UK) for 5 minutes at 37°C/5% CO_2_. Experiments involving the treatment of CLL cells with interleukin 4 (IL4) were performed as outlined previously.^22^ Briefly, following recovery, counting and dilution, cells were either treated with 10 ng/ml IL4 (R&D Systems, Abingdon, UK) or left untreated and incubated for 24 hours at 37°C/5%CO_2_. The following day, IL4-treated cells were stimulated with 20 µg/ml anti-IgM F(ab’)_2_, while untreated cells were treated with anti-IgM or isotype control F(ab’)2 (negative control). MEC-1 cells were pre-treated and incubated at 37°C/5%CO_2_ for 1 hour with 500nM ibrutinib or dasatinib prior to protein lysis.

### Protein lysis for kinobead analysis

Cells were washed twice using ice-cold phosphate-buffered saline and resuspended in kinobead lysis buffer (50 mM HEPES (pH 7.5), 150 mM NaCl, 0.5% (v/v) Triton X100, 1 mM EDTA, 1 mM EGTA, 10 mM NaF, 2.5 mM NaVO_4_, 1x Protease Inhibitor Cocktail (Roche, Sussex, UK), and 1x Phosphatase Inhibitor Cocktails 2 and 3 (Sigma, Dorset, UK)). Suspensions were incubated on ice for 10 minutes with regular mixing before sonication and clarification by centrifugation at 13000 rpm for 10 minutes. The supernatant was collected and protein content quantified using BCA assay (Thermo Fisher Scientific). Lysates were stored at -80 °C prior to analysis.

### Kinobead preparation

We employed a protocol similar to that previously described.^12^ The broad-spectrum kinase inhibitor Ki-NET (CTx-0294885) was conjugated to NHS-activated Sepharose4 Fastflow resin (GE Healthcare, Little Chalfont, UK). Briefly, Ki-NET was solubilized in coupling buffer consisting of 50% dimethylformamide/50% sodium phosphate (pH 7.0). Sepharose was resuspended and transferred onto the filter of a 0.22 µM pore filter flask (Corning). The filter was then washed twice with 0.5 M NaCl, once with Millipore water and once using coupling buffer. The washed beads were recovered from the filter and added to the dissolved Ki-NET. To promote conjugation, 1 M *N*- (3Dimethylaminopropyl)-N’-ethylcarbodiimide hydrochloride (EDC) was added and KiNET beads were incubated overnight at room temperature in the dark with continuous end-on-end mixing. The following day, beads were washed twice and resuspended in coupling buffer containing 1 M ethanolamine. EDC (1 M) EDC was added and the beads were incubated overnight as described above. Conjugated kinobeads were washed three times with coupling buffer, twice with Millipore water and once with 20% (v/v) ethanol before final resuspension in 20% ethanol and being stored at 4°C. For other kinobeads tested, the above process was used for the inhibitor VI16832 and purvalanol-B which were conjugated to EAH Sepharose, while bisindolylmaleimide-X (Bis-X), dianillopyrimidine and pp58 were conjugated to a 1:1 mixture of NHSactivated Sepharose-4 Fastflow resin and EAH Sepharose.

### Isolation of kinases using kinobeads

Protein lysates were thawed on ice and the specific quantity required (3 mg for MEC1 experiments, 1 mg for primary CLL) was aliquoted, diluted with kinome lysis buffer and adjusted to 1 M NaCl. Polyprep columns (BioRad) were placed in a rack and kinobeads were resuspended, applied carefully to the bottom of the column and washed using a high salt buffer (50 mM HEPES (pH 7.0), 1 M NaCl, 0.5% (v/v) Triton X-100, 1 mM EDTA, 1 mM EGTA). In a separate column, unconjugated Sepharose was added to create a block condition and washed using high salt buffer. Block columns were placed above the kinobead columns and lysates were carefully loaded. Once lysates had passed through both the block and kinobead columns, the block columns were discarded and the kinobead columns were washed using high salt buffer, then low salt buffer (50 mM HEPES (pH 7.0), 150 mM NaCl, 0.5% (v/v) Triton X-100, 1 mM EDTA, 1 mM EGTA) and then SDS wash buffer (low salt buffer supplemented with 0.1% (w/v) SDS). Columns were stoppered to prevent flow-through before the addition of elution buffer (0.5% (w/v) SDS, 1% (v/v) b-mercaptoethanol, 0.1 M Tris-HCl (pH 6.8) in LC-MS grade water). Columns were capped and incubated at 98 °C for 15 minutes before being collected into low-bind Eppendorf tubes.

### Processing of kinobead elutions

Elutions from kinobeads were initially reduced by adding 5 mM DTT and incubating at 60 °C for 25 minutes. Samples were alkylated with 20 mM iodoacetamide (IAA) and incubated at room temperature for 30 minutes in darkness. Alkylation was inhibited by adding another 5 mM DTT and incubation at room temperature for 5 minutes in darkness. Samples were loaded into concentrator spin-columns (Millipore, Herts, UK) and centrifuged at 3000 rpm for 30 minutes at 4 °C. Concentrated samples were collected into low-bind Eppendorf tubes. To precipitate proteins, samples underwent initial spin-washes using methanol, chloroform and water (all LC-MS grade) before centrifugation at 13,000 rpm for 10 minutes at 4 °C. The interphase layer (protein) was collected and washed using MS-grade methanol. Samples were dried by speedvac before resuspension in 50 mM HEPES (pH 8.0) and the addition of 500 ng Trypsin Gold (Promega, Southampton, UK). Pellets were incubated overnight at 37 °C. The following day, peptide samples were washed 3 times using water-saturated ethyl acetate to remove Triton X-100 before drying with a speedvac. Peptides underwent de-salting using C-18 spin columns (Thermo Fisher Scientific). Columns were equilibrated using a buffer of 5% acetonitrile (ACN)/0.5% trifluoroacetic acid (TFA). Sample pellets were resuspended in this buffer and loaded into prepared spin columns and centrifugation at 4,400 rpm for 1 minute. Columns underwent further washed using equilibration buffer before undergoing elution using 50% ACN and stored at 80°C prior to mass spectrometry (MS).

### MS sample testing and data analysis

Peptide samples were supplied to the WPH Proteomics Facility of the University of Warwick where they underwent final preparation and MS analysis using an Orbitrap Fusion instrument (Thermo Fisher Scientific) as outlined previously.^23^ Data files generated by MS (RAW) were analyzed using MaxQuant software (v1.5.3.8; Max Planck Institute). Peptide intensities involving serine, threonine, tyrosine phosphorylation (PSTY) alongside oxidation and acetylation were used as variable modifications, while carbamidomethylation to cysteine was using as a fixed modification. Protein identification was made by comparison to a Uniprot human proteome database. A 1% false discovery rate (FDR) was used for all searches. Changes in kinase levels were determined by creating ratios comparing either sIgMstimulated CLL cells to the relevant control antibody-treated cells for each patient. Directionality of response was achieved by converting ratios to log2 scale. These data were then used to create heatmaps using Multi-experiment Viewer (MeV) software (tm4.org).

### Immunoblotting

For direct analysis, cells were lysed using radioimmunoprecipitation assay (RIPA) buffer whereas for kinobead analysis, elutions from kinobeads were mixed with and equal volume of 2x loading buffer prior to SDS-PAGE. Immunoblotting was performed using anti-phospho-ERK1/2 (Y202/T204), anti-ERK, anti-phospho-AKT (S473), antiAKT or anti-BTK (all Cell Signaling Technologies, Hitchin, UK). Anti-actin (Sigma) was used as a loading control. Bound primary antibodies were detected using speciesspecific fluorophore-conjugated secondary antibodies (LiCOR Biosciences, Cambridge, UK) and imaged using a LiCOR Odyssey CLx instrument.

### Nanostring kinase mRNA analysis

RNA was extracted using an RNA-Easy kit (Qiagen) in accordance with the manufacturer’s protocol. RNA quantity and quality were assessed using a Nanodrop 1000 instrument (Thermo Fisher Scientific). Kinase gene analysis was carried out using nCounter XT HuV2 Kinase arrays (Nanostring, Seattle, WA, USA) according to the manufacturer’s protocol. 100 ng of RNA was ligated to capture and reporter probes by incubation at 65°C for 22 hours using a Veriti Thermal Cycler (Thermo Fisher Scientific). Preparation of array cartridges was performed using a Nanostring PrepStation and read using an nCounter reader. Data were analyzed using nSolver software (v3.0; Nanostring).

### Phosphatase analysis

For surface expression analysis, cells were prepared as outlined above and 1×10^6^ were aliquoted into FACS tubes. Cells were pelleted by centrifugation and each tube was resuspended in flow buffer (1% (w/v) bovine serum albumin, 4 mM EDTA and 0.15 mM NaN_3_ in phosphate-buffered saline) containing anti-CD19 and anti-CD5 antibodies for CLL cell detection. Antibodies against specific key markers and surface phosphatases were then used in various combinations. Cells were incubated on ice in the dark for 15 minutes prior to washing with flow buffer. Cells were resuspended in flow buffer and analyzed using a Canto cytometer. Data were analyzed using FlowJo.

For intracellular phosphatases, lysates were analyzed by immunoblotting with the following antibodies: anti-phospho-SHIP1, anti-phospho-SHP1, anti-phospho-LYN (Y507) (all Cell Signaling Technologies), anti-SHP1, anti-phospho-LYN (Y396) (both Abcam, Cambridge, UK), anti-SHIP1, anti-LYN (both Insight Biotechnology, Middlesex, UK), anti-phospho-PTPN22 (R&D Systems) and anti-GAPDH (Thermo Fisher Scientific). Primary antibodies were probed using species-specific HRPconjugated secondary antibodies (Dako) and viewed using a Chemidoc imager (BioRad).

### Data sharing

All kinome signatures detected by Nanostring and kinobead-MS are listed in **Supplementary List**. For data values, please contact.

## Acknowledgments

The authors would like to gratefully thank Drs Melanie Oates, Ke Lin, Alix Bee, Gillian Johnson and Jessica Bithell (Molecular and Clinical Cancer Medicine, University of Liverpool) as well as Dr Lindsay Smith (University of Southampton) for providing clinical information and aiding in sample retrieval. In addition, we would like to thank Drs Cleidiane Zampronio and Juan Hernandez-Fernaud of the WPH Proteomics Facility (University of Warwick) for performing MS sample analysis and Drs Lucille Rainbow and Pia Koldkjaer of the Centre for Genomic Research (CGR), University of Liverpool for their assistance in Nanostring analysis. Moreover, we would like to thank Drs Rebecca Jarvis and Sophie Cramp from the Cancer Research UK Clinical Trials Unit, University of Birmingham, for providing information pertinent to the IcICLLe trial samples. Finally, and most importantly, we would like to humbly thank the patients who provided samples for the work performed in this study.

## Authorship

AJL optimized, designed and performed experiments, analyzed data, wrote, reviewed edited the manuscript and obtained funding. LIK, AM and AD performed experiments and interpreted data. NK, ARP and FFF supplied clinical samples/input and reviewed the manuscript. AR and PH provided data and input concerning the clinical trial samples. DJM designed experiments and reviewed manuscript and obtained funding. AJS, GP, IAP and JRS designed experiments, interpreted data, reviewed and edited the manuscript and obtained funding.

## Supplementary Figure Legends

**<u>Supplementary Figure 1</u>**: **A**; Immunoblot analysis of MEC-1 cells showing expression of total and phospho-ERK-1/2 and AKT in control (NA), ibrutinib or dasatinib-pre-treated cells (both at 500 nM for 60 minutes). Actin was analyzed as an additional loading control. **B**; Venn diagrams showing overlap of kinases isolated by the different KI used to create kinobeads during this study. **C**; Correlation graphs comparing intensities for kinases isolated from MEC1 lysates by Ki-NET beads in untreated, ibrutinib-and dasatinib pre-treated cells.

**<u>Supplementary Figure 2:</u> A**; Graph comparing kinases identified at either mRNA level, protein (by kinobead isolation) or both in our patient cohort. **B**; Correlation graphs for 2 representative CLL patients, comparing intensities gained for our kinome signature for 2 biological repeats for baseline (Control F(ab’)2) cells and in response to anti-IgM treatment. **C**; Heatmaps showing kinome fingerprints for CLL patients stratified according to *IGHV* mutation status, Binet staging or karyotype status. **D**; Graphs illustrating relative change in isolation of SYK and BTK in primary CLL cells from patients stratified according to *in vitro* iCa^2+^ flux response as being Non-Signaler (NS; iCa^2^<5%), or Signaler (S; iCa^2+^>5%). **E**; Correlation of kinase mRNA expression determined by Nanostring between Treatment Naïve (TN) and Previously Treated (PT) patients.

**<u>Supplementary Figure 3:</u> A**; Flow cytometric analysis of inhibitory coreceptors and **B;** immunoblot analysis of phosphatases in three samples from untreated (UT) patients and three samples from previously treated (PT) patients. In **A**, graph shows results for individual samples and mean (±error). In **B**, GAPDH was analyzed as an additional loading control.

**<u>Supplementary Figure 4:</u> A**; Relative changes to surface IgM (sIgM) (GeoMFI) expression for 4 IcICLLe trial patients between baseline and 1 month after initial receipt of ibrutinib treatment. **B**; Immunoblotting of to illustrate loss of BTK binding to kinobeads through in vivo action of ibrutinib. Kinobead isolation was performed on matched samples for 2 representative patients, comparing the pre-ibrutinib sample (TP1) to a sample taken following receipt of ibrutinib (TP2). Kinobead elutions were mixed with loading buffer and separated using SDS-PAGE prior to blotting and probing.

## Notes

The authors declare no conflict of interest.

### Competing Interest Statement

The authors have declared no competing interest.

