## Supplementary figures and images for "Kinobead Profiling Reveals Reprogramming of B-cell Receptor Signaling in Response to Therapy Within Primary Chronic Lymphocytic Leukemia Cells"

### Supplementary Figure 1

#### Supplementary Figure 1

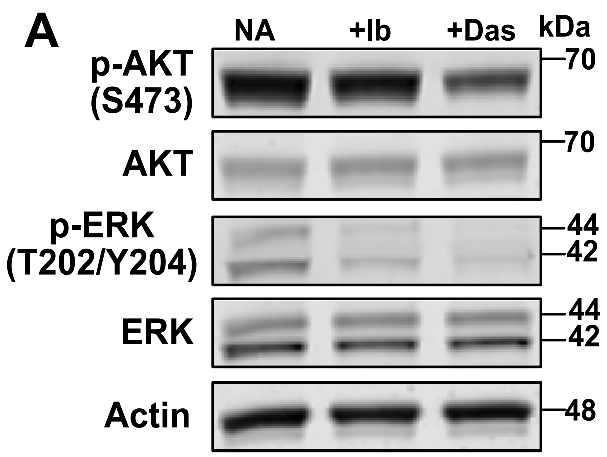


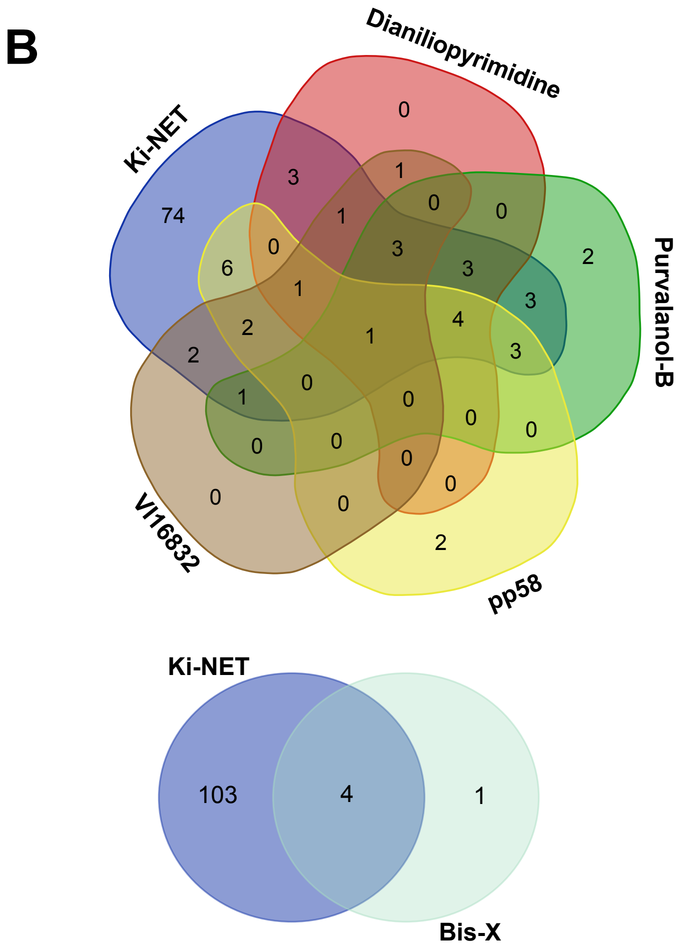


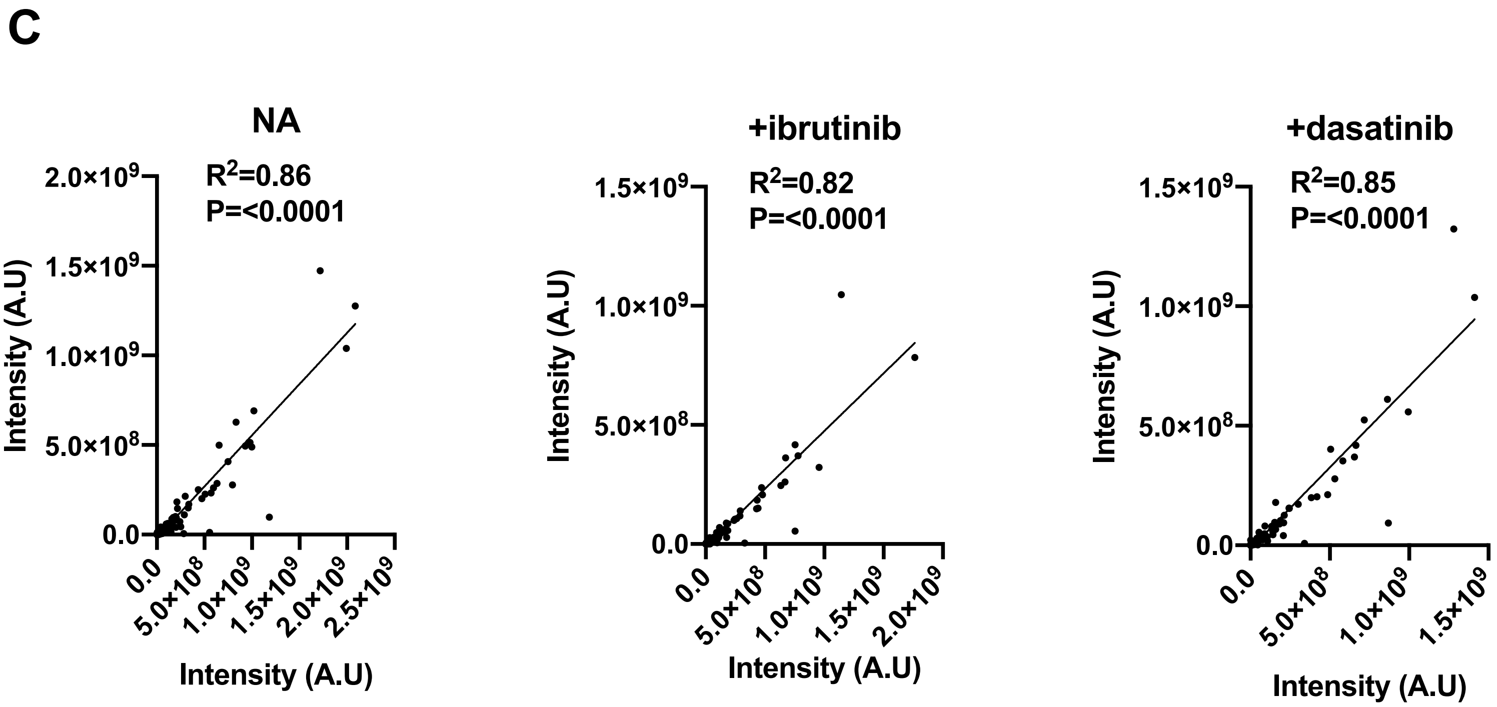

### Supplementary Figure 2

#### Supplementary Figure 2

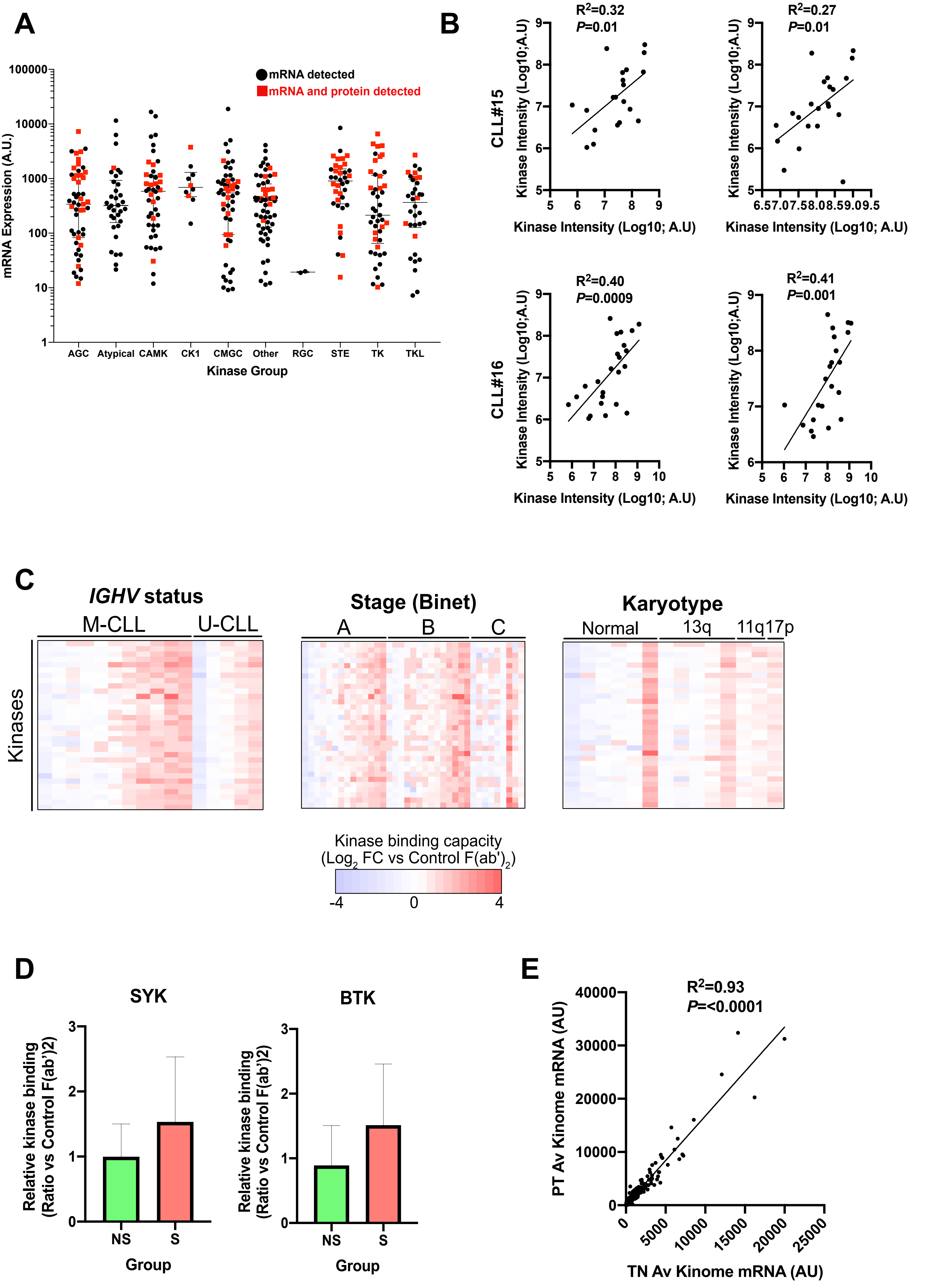

### Supplementary Figure 3

##### Supplementary Figure 3

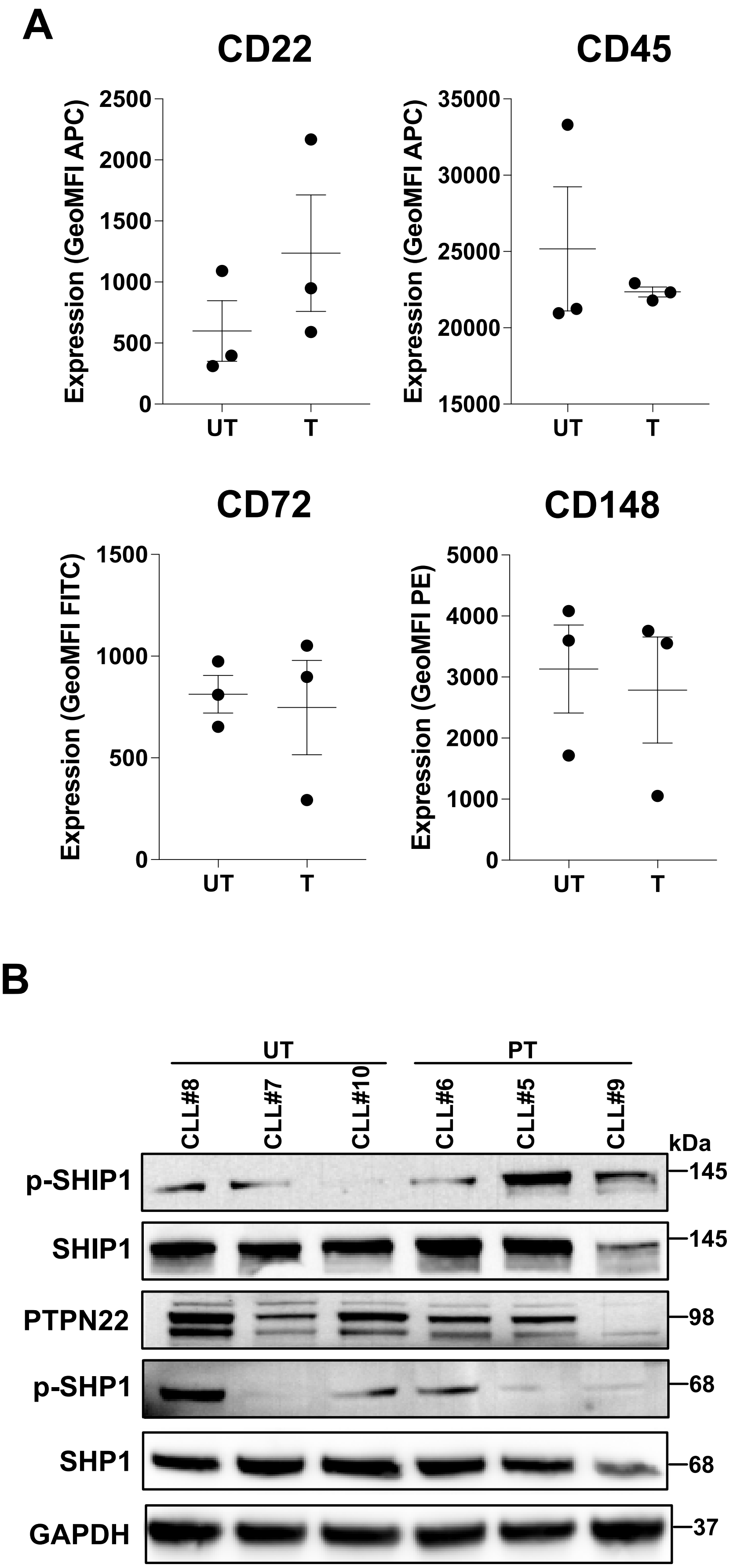

### Supplementary Figure 4

#### SUPPLEMENTARY FIGURE 4

**A**

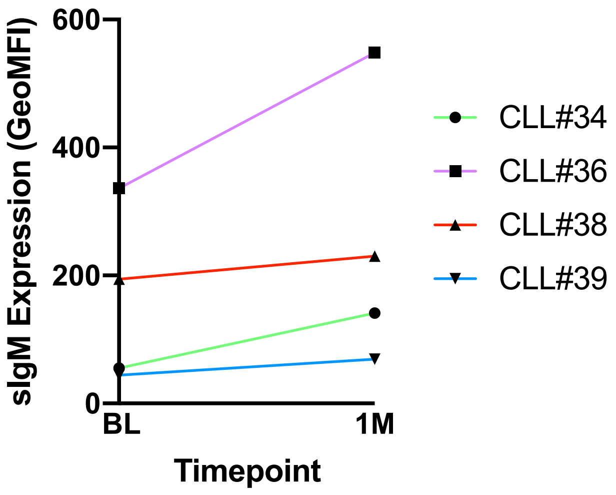


**B**

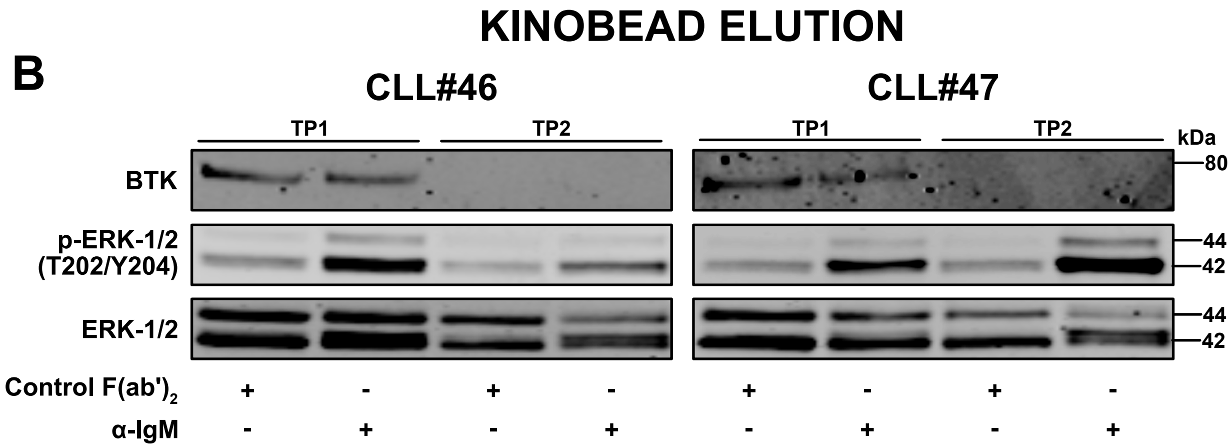
